# Population-scale detection of non-reference sequence variants using colored de Bruijn Graphs

**DOI:** 10.1101/2021.03.23.436560

**Authors:** Thomas Krannich, W. Timothy J. White, Sebastian Niehus, Guillaume Holley, Bjarni V. Halldórsson, Birte Kehr

## Abstract

**Motivation:** With the increasing throughput of sequencing technologies, structural variant (SV) detection has become possible across tens of thousands of genomes. Non-reference sequence (NRS) variants have drawn less attention compared to other types of SVs due to the computational complexity of detecting them. When using short-read data, the detection of NRS variants inevitably involves a *de novo* assembly which requires high-quality sequence data at high coverage. Previous studies have demonstrated how sequence data of multiple genomes can be combined for the reliable detection of NRS variants. However, the algorithms proposed in these studies have limited scalability to larger sets of genomes.

**Results:** We introduce *PopIns2*, a tool to discover and characterize NRS variants in many genomes, which scales to considerably larger numbers of genomes than its predecessor *PopIns*. In this article, we briefly outline the PopIns2 workflow and highlight our novel algorithmic contributions. We developed an entirely new approach for merging contig assemblies of unaligned reads from many genomes into a single set of NRS using a colored de Bruijn graph. Our tests on simulated data indicate that the new merging algorithm ranks among the best approaches in terms of quality and reliability and that PopIns2 shows the best precision for a growing number of genomes processed. Results on the Polaris Diversity Cohort and a set of 1000 Icelandic human genomes demonstrate unmatched scalability for the application on population-scale datasets.

**Availability:** The source code of *PopIns2* is available from https://github.com/kehrlab/PopIns2.

## 1 Introduction

The genome of every person contains sequence that is not present in the current reference genome [Faber-Hammond and Brown, 2016]. This additional sequence differs between people and contributes to genetic diversity. It can disrupt exons or regulatory elements [Wong *et al.*, 2020] and associate with diseases [Kehr *et al.*, 2017] analogous to single-nucleotide variants, small insertion-deletion (indel) variants and larger copy-number, inversion and translocation variants. The vast majority of sequence missing in the reference genome is repetitive [Delage *et al.*, 2020, Manni and Zdobnov, 2020]. Owing to collaborative efforts such as the Telomere-to-Telomere (T2T) consortium [Miga *et al.*, 2020, Logsdon *et al.*, 2021] we can expect that the major sequence content of even highly repetitive centromere and telomere regions in the human genome will be known soon. Possibly we will discover variants of highly repetitive regions in the future, for example copy number variants, but the specific sequence of consecutive base pairs that constitute repeat regions will be near complete.

A smaller portion of sequence missing from the reference genome is present only in a subset of the population, the NRS variants. Others refer to NRS variants as insertions [Wong *et al.*, 2020, Delage *et al.*, 2020] because the variants describe novel sequence with respect to the reference genome. However, the majority of NRS appears to be ancestral rather than novel because they can be found in other primate genomes [Lee *et al.*, 2020, Kehr *et al.*, 2017]. A convincing explanation for the existence of NRS is that the genomes used to construct the reference genome lacked these sequences. In light of this, it may be misleading to refer to NRS variants as insertions or novel sequences.

Compared to other types of structural variation, NRS variants are less well studied because their detection is algorithmically more challenging. While deletions, duplications, inversions and translocations can be characterized by breakpoints in the reference sequence, NRS variant detection additionally involves determining unknown sequence. If the NRS is longer than the read length, a sequence assembly is inevitable. Sequence reads consisting only of NRS do not align to the reference genome and, therefore, they do not indicate the candidate variant positions. As a result, NRS variant detection comprises assembling unaligned reads into candidate NRS, determining the positions of NRS variants in the reference genome based on partially aligned reads, and genotyping the detected NRS variants.

Early NRS detection methods, such as Pindel [Ye *et al.*, 2009] or SOAPindel [Li *et al.*, 2013], relied on the alignment of one read in a read pair. They were therefore limited to short NRS variants. The first methods that included fully unaligned read pairs required substantial computational resources and were very limited in scalability. For example, Cortex [Iqbal *et al.*, 2012] introduced the colored de Bruijn graph (CDBG) data structure to assemble several genomes simultaneously. In the resulting *de novo* whole genome assemblies, NRS variants can be traced just like any other type of structural variant. The tool Cortex itself is very limited in terms of scalability and cannot handle more that a dozen genomes simultaneously. As a consequence, the scientific community has yielded considerably improved implementations of the CDBG data structure [Muggli *et al.*, 2019, Wittler, 2020, Almodaresi *et al.*, 2018] since. Similarly, the tools MindTheGap [Rizk *et al.*, 2014] and Basil&Anise [Holtgrewe *et al.*, 2015] can only handle a single genome. Both approaches first identify candidate variant end positions in the read alignment and implement custom procedures to assemble the NRS between two NRS variant end positions.

While the mentioned approaches present algorithmically interesting solutions to the NRS detection problem, they do not scale to large data sets. Only the development of data-focused NRS variant calling pipelines more recently enabled the analysis of hundreds, thousands or even tens of thousands of genomes. Early pipelines determined NRS contigs without their positions in the reference genome, for example in 769 Dutch genomes [Hehir-Kwa *et al.*, 2016], 10,000 genomes of several ancestries [Telenti *et al.*, 2016], and 300 genomes from 142 diverse populations [Mallick *et al.*, 2016]. More recently, studies of NRS variants include precise breakpoint positions and genotype estimates, for example for 15,219 Icelanders [Kehr *et al.*, 2017] and 910 African genomes [Sherman *et al.*, 2019]. Some pipelines for moderate numbers of genomes create whole-genome assemblies prior to NRS variant calling, such as the pipelines that were applied to 50 Danish trios [Maretty *et al.*, 2017, Liu *et al.*, 2015], 275 Han Chinese genomes [Duan *et al.*, 2019], 1000 Swedish genomes [Eisfeldt *et al.*, 2020], and 338 genomes from genetically divergent human populations [Wong *et al.*, 2018, Wong *et al.*, 2020]. Finally, pipelines developed for the 1000 genomes project data [Lee *et al.*, 2020] and for the TOPMed program [Taliun *et al.*, 2021] search for NRS variants that match related genomes like other primates’ genomes.

In contrast to the earlier algorithmic approaches for NRS variant calling, many of the pipelines for large numbers of genomes have not been benchmarked and released as stand-alone tools for application to other data sets. One exception is our tool PopIns [Kehr *et al.*, 2016], which we used to analyze the Icelandic genomes [Kehr *et al.*, 2017]. PopIns assembles unaligned reads into contig sequences per genome, merges the contig sequences from all input genomes into a single set of NRS, and determines positions and genotypes of NRS variants. By merging contigs sequences across genomes, PopIns takes advantage of the increased total coverage of NRS variants that occur in more than one genome. In addition, combining data from all genomes early in the process, eliminates the need to later heuristically combine variant calls from different individuals.

Since its publication, PopIns has been challenged by new NRS detection tools, most importantly by Pamir [Kavak *et al.*, 2017]. Pamir works well on small numbers of genomes and was shown to have superior accuracy compared to PopIns on simulated data. This led us to investigate weaknesses of PopIns, resulting in the merging step being identified as responsible for both false positive and false negative NRS variant calls, as well as for limiting the number of genomes due to high memory requirements.

PopIns’ merging step combines contigs assembled from unaligned reads of many genomes into a single, preferably non-redundant set of NRS. We previously noted its similarity to a classical genome assembly problem [Kehr *et al.*, 2016]. Classical genome assembly for short read data is commonly based on de Bruijn graphs (DBG) [Pevzner *et al.*, 2001, Compeau *et al.*, 2011]. The tool Cortex [Iqbal *et al.*, 2012] augmented DBGs with colors in order to simultaneously process several genomes. Recently, a number of elaborate, highly space-efficient and versatile implementations of CDBG have been introduced [Almodaresi *et al.*, 2017, Almodaresi *et al.*, 2018, Muggli *et al.*, 2017, Muggli *et al.*, 2019, Mustafa *et al.*, 2019, Holley and Melsted, 2020, Karasikov *et al.*, 2020, Khan and Patro, 2021, Alanko *et al.*, 2021]. One such implementation, the Bifrost API [Holley and Melsted, 2020], is used in this work.

In this paper, we introduce an NRS detection approach that reaches high accuracy on simulated data and scales to many genomes in practice. Specifically, we describe an alternative merging step for PopIns, which models the task as a weighted minimum path cover problem on a CDBG. Our heuristic solution to this NP-hard problem defines novel rules for traversing the CDBG according to path coverage and color information. We implemented the approach in PopIns2 together with minor improvements in the other steps of the original PopIns tool and show that PopIns2 outperforms PopIns and matches the accuracy of Pamir on simulated data. Requiring orders of magnitude less memory for the merging step, the scalability of PopIns2 exceeds that of PopIns and Pamir by far.

## 2 Methods

While PopIns2 introduces substantial improvements compared to PopIns, both tools follow the same overall workflow as previously described [Kehr *et al.*, 2016, Kehr *et al.*, 2017]. In the following, we first summarize the individual steps in this workflow (Supplementary Figure 1), highlighting differences between PopIns and PopIns2. Subsequently, we describe the newly developed algorithm for the merging step in detail.

### 2.1 Overview of the PopIns2 workflow

Just like PopIns, PopIns2 takes as input a collection of short read sequences from many genomes, which have been aligned to a reference genome, in the BAM file format. The final output is a set of NRS and their positions in the reference genome together with genotypes and genotype likelihoods for all genomes. The NRS are given in the FASTA format. Precise positions may be given for one or both endpoints of the sequences. We specify positions in the breakend notation of the VCF file format.

#### Assembly step

The assembly step processes a single genome at a time. It selects all reads that do not align or align only poorly to the reference genome and passes them to a classical genome assembly tool to compute contiguous sequences (contigs). For the process of selecting poorly aligned reads, we identified the alignment score factor (ASF) parameter of PopIns to have a large impact on the contig assembly. The parameter determines a threshold for the ratio between the read alignment score (AS tag) and the read length, when filtering for poorly aligned reads. In PopIns2, we implemented the ASF as a command line parameter (default 0.67) but kept all other parameters as previously described [Kehr *et al.*, 2017].

During the development of the original PopIns module, we assessed several genome assembly tools [Zimin *et al.*, 2013, Bankevich *et al.*, 2012, Zerbino and Birney, 2008] for integration into the workflow. For PopIns2, we compared the Velvet assembler [Zerbino and Birney, 2008] used in PopIns to the more recent Minia assembler [Chikhi and Rizk, 2013] in terms of precision and recall of the subsequent merge module. Based on an improved precision and F1-score (Supplementary Table 2), we changed the default internal assembly tool in PopIns2 to Minia but maintain compatibility with Velvet.

#### Merging step

The merging step takes as input contigs output by the assembly step from many genomes and combines them into a single set of candidate NRS, which we previously termed supercontigs. The merging step from PopIns has been replaced by an entirely new merging approach in PopIns2. The new approach constructs a CDBG from the input contigs and we formulate the problem of finding candidate NRS as a minimum path cover problem on the CDBG. Below we suggest a heuristic algorithm that traverses a CDBG and uses the colors to identify a set of candidate NRS.

#### Placing step

The placing step determines potential positions of the NRS in the reference genome. It underwent no changes in PopIns2 compared to the latest version of PopIns; details have been described previously [Kehr *et al.*, 2017]. Briefly, we utilize the information of read pairs, where one read is aligned to the reference genome and the other end is not. For each such read pair, an alignment to an NRS is found for the unmapped read. Its corresponding reference-aligned read provides candidate insertion positions (intervals) and determines the orientation of the NRS. Some assembled sequences comprise flanking reference sequences of considerable length in addition to the NRS. In these cases, PopIns split aligns contig ends to the interval in the reference genome for determining exact insertion positions of those NRS. For many remaining NRS, split alignments of reads provide exact predictions of the insertion positions at base pair resolution.

#### Genotyping step

The final genotyping module of PopIns2 computes genotypes and genotype likelihoods for every NRS variant in every individual using the NRS, its position in the reference genome and the individual’s set 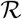 of sequencing reads. Details about this module have been described previously [Kehr *et al.*, 2016, Kehr *et al.*, 2017] and underwent no algorithmic changes in PopIns2 since the latest version of PopIns. Briefly, the algorithm constructs the sequence of the alternative allele with the NRS inserted and extracts the reference sequence without the NRS around an insertion position (reference allele), aligns the reads in 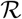 to both alleles and computes the likelihoods for an NRS to be present on 0, 1 or 2 of the individual’s chromosomal copies.

### 2.2 Merging NRS of many genomes using a CDBG

This section describes the new merging step of PopIns2 starting with all necessary definitions, followed by the problem formulation and, finally, our heuristic solution.

#### 2.2.1 Definitions

##### Genomic sequences

Let *s* ∈ Σ^*m*^ be a genomic sequence of length *m* over the DNA alphabet Σ = {*A, C, G, T*}. We denote the reverse complement of *s* as 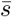. A substring of *s* is defined by a starting position and a length *l*. A prefix of *s* is a substring that starts at the first position of *s* and a suffix is a substring that ends at the last position of *s*. Because a prefix (suffix) of *s* is defined by its length *l*, we refer to it as an *l*-prefix (*l*-suffix) of *s*. A *k*-mer is a sequence of length *k*.

##### Directed graphs

In a directed graph *G* = (*V, E*), which consists of a set of vertices *V* and a set of edges *E* = *V* × *V* , an edge *e* is an ordered pair (*u, v*) of vertices *u, v* ∈ *V*. We call *v* a successor of *u* and *u* a predecessor of *v*. The in-degree (out-degree) of a vertex *v* denotes the number of its predecessors (successors). A walk *p* = *v*_1_*, … , v_n_* through the graph is a sequence of vertices such that *v*_*i* + 1_ is a successor of *v*_*i*_ for 1 ≤ *i* < *n*. A path is a walk without circuits, i.e., in which all vertices are distinct. A path *p* = *v*_1_*, …, v_n_* is non-branching if vertex *v*_1_ has out-degree 1, vertex *v_n_* has in-degree 1 and all other vertices *v*_2_ , …, *v_n_* – 1 have both in-degree 1 and out-degree 1. A non-branching path *p* = *v*_1_ , …, *v_n_* is maximal if the in-degree of *v*_1_ and the out-degree of *v_n_* are both unequal to 1. Let *V_p_* be the set of vertices forming the path *p*. A set of paths *P* = {*p*_1_*, p*_2_*, …, p_m_*} for graph *G* = (*V, E*) forms a path cover of *G* if each vertex *v* ∈ *V* is part of at least one path in *P*, i.e., *V* = ⋃_*p*∈*P*_ *V*_*p*_.

##### De Bruijn graphs

A de Bruijn graph (DBG) for a given *k* over a set of input sequences *S* is a directed graph *G* = (*V, E*), such that each *k*-mer that is a substring of either *s* or 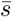 with *s* ∈ *S* corresponds to a vertex *v* ∈ *V*. An edge *e* = (*u, v*) ∈ *E* exists if the (*k* – 1)-suffix of the *k*-mer corresponding to *u* equals the (*k* – 1)-prefix of the *k*-mer corresponding to *v*. A walk or path *p* of *n* vertices in a DBG corresponds to a genomic sequence *ω*(*p*) of length *n* + *k* − 1. Genomic sequences that correspond to maximal non-branching paths in a DBG are called unitigs.

In a compacted DBG *G*′ = (*V* ′, *E*′), all maximal non-branching paths of the original DBG are represented by a single vertex. Thus, vertices *u, v* ∈ *V* ′ correspond to unitigs and an edge *e* = (*u*, *v*) ∈ *E*′ exists if the (*k* – 1)-suffix of the unitig corresponding to *u* equals the (*k* − 1)-prefix of the unitig corresponding to *v*. A maximal unitig path *p* = *v*_1_, *v*_2_, *…, v_m_* is a path through a compacted DBG where the in-degree of *v*_1_ and the out-degree of *v_n_* are both equal to 0.

To better represent double-stranded DNA, many implementations of DBGs are bidirected. In a bidirected DBG a single vertex represents both a *k*-mer (or unitig) and its reverse complement. Furthermore, edges have two types of orientation [Medvedev *et al.*, 2007]. That is, edges connect to either orientation of the *k*-mer (unitig). For simplicity and clarity, we describe our method using simple DBGs below while our implementation is based on the Bifrost API [Holley and Melsted, 2020], which uses bidirected DBGs.

##### Colored de Bruijn graphs

While DBGs represent a single sequence set, a colored de Bruijn graph (CDBG) can represent a set of several sequence sets. In a CDBG *G* = (*V, E, C*) over a set of sequence sets *S* = {*S*_1_, *S*_2_, …, *S_n_*}, vertices are additionally labeled with bitvectors of length *n* that store information about the presence of *k*-mers in each sequence set *S_i_* ∈ *S*. Each bit in the bitvector represents a sequence set *S_i_* ∈ *S* and is thought of as a color. The *i*’th bit in the bitvector is set to 1 if and only if the corresponding *k*-mer occurs at least once in *S_i_*. In a colored compacted DBG, a vertex corresponding to a unitig of length *l* is labeled with *l − k* + 1 bitvectors, one per *k*-mer of the unitig. Typically, the bitvectors of all *k*-mers are stored in a matrix *C* = (*c_ij_*), where each entry *c_ij_* indicates the presence of *k*-mer *j* in sequence set *S_i_*. The matrix is called the color matrix of the (compacted) CDBG.

##### Jaccard index

The Jaccard index is a measure for the similarity of sets [Jaccard, 1912]. It is defined as the ratio of the intersection and the union of two sets. Analogously, the definition of the Jaccard index on two bitvectors *x* and *y* is

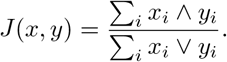

#### 2.2.2 Problem formulation

Given the sets of contigs created in the assembly step for all input genomes, our goal in the new merging step is to extract NRS from a compacted CDBG built from the set of contig sets. While whole genome assembly aims to minimize the number of contig sequences and maximize contiguity, we are looking for many comparably short genomic sequences. NRS assembly is in fact more related to transcript assembly [Xing *et al.*, 2004, Trapnell *et al.*, 2010] than whole-genome assembly with the difference that transcript assembly typically operates on a relatively small DAG and we are given a CDBG with cycles. Similar to transcript assembly, we observe that the paths corresponding to the sought NRS form a path cover of the CDBG when assuming that all *k*-mers in the CDBG are part of at least one NRS in the solution. A naive solution for finding a path cover is to enumerate all possible paths in the graph. This naive solution, however, leads to unwanted redundancy and many of the paths will not correspond to actual NRS.

Therefore, we formulate our problem as a minimum path cover problem, which is to find the smallest number of paths that form a path cover in a given graph. This problem is NP-complete [Garey and Johnson, 1990] on graphs other than directed acyclic graphs (DAGs) [Lawler, 2001], since determining whether a single path suffices is equivalent to determining whether the graph has a Hamiltonian path [Rizzi *et al.*, 2014]. Note that some of the vertices can be part of several paths in a minimal path cover, which corresponds to shared sequences in biology, e.g., mobile elements.

We further extend the problem formulation by the color information. Our idea is that all *k*-mers along a path that corresponds to an actual NRS should be labeled with a similar set of colors, i.e. with those colors corresponding to the genomes that carry the NRS. We define a color weight function *ϕ* : *p* ⟶ ℝ that assigns a weight to each path *p* = *u*_1_, …, *u*_*n*_ through a compacted CDBG as

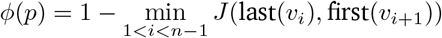

where first(*v*) and last(*v*) are the bitvectors corresponding to the first and last *k*-mer of vertex *v*. Using this weight function the weighted minimum path cover problem is to find a path cover *P* = {*p*_1_, *…, p_n_*} of *G* that minimizes the weight Σ_*p*∈*P*_ *ϕ*(*p*). An optimal solution to the weighted minimum path cover problem then corresponds to a set of candidate NRS merged from contig sets of many genomes.

#### 2.2.3 A greedy heuristic

As a practical solution, we suggest a greedy heuristic to solve the weighted minimum path cover problem on a compacted CDBG built from contig sets *S_i_* assembled from unaligned and poorly aligned reads of many genomes. The key idea is to start a greedy depth first search (DFS) from the sources of the graph, i.e., vertices that have only successors but no predecessors. During the DFS traversal, vertices are prioritized based on the Jaccard index of color vectors. The recursion of the traversal continues until the search reaches a sink, i.e, a vertex without successors, or aborts due to local substructures that cannot be resolved. In case it reaches a sink, the sequence of vertices is returned if the sequence comprises a minimum number of not yet covered *k*-mers.

More formally, let *G* = (*V, E, C*) be a compacted CDBG for the set of sequence sets *S* = {*S*_1_, *S*_2_, *…, S_n_*} and the positive integer *k*. We label each vertex (unitig) *u* ∈ *V* with a traversal state which is either seen or unseen. Initially all vertices are unseen. In addition, we store a set 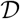 of covered vertices that is initially empty. Note that each vertex *u* can have at most four predecessors and at most four successors since we use the DNA alphabet.

In the initialization step we check every vertex *u* ∈ *V* for the presence of predecessors or successors. We distinguish three possible outcomes of this check: i) *u* has no predecessors and no successors (*u* is a singleton), ii) *u* has predecessors and iii) *u* has only successors (*u* is a source). If *u* is a singleton, we return its sequence and continue with the next vertex. If *u* has predecessors, we just continue with the next vertex. If *u* is a source, we pass it to our recursion step.

The recursion step marks the current vertex *u_c_* as seen and checks whether there are unseen successors. If the current vertex *u_c_* is a *sink*, we count the number of *k*-mers along path *p* from source to sink that are not part of a vertex in 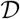. If *p* contains more than *τ* novel *k*-mers we add all vertices in *p* to 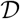 and return the NRS *ω*(*p*).

If *u_c_* has a non-empty set of unseen successors 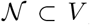, we need to decide how to continue the traversal of the CDBG. This decision is a crucial design choice of the algorithm since the traversal order of the vertices determines the resulting contigs. We here utilize the DBG colors. We compute a weight for each edge from *u_c_* to each successor 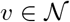 using the bitvectors last(*u_c_*) corresponding to the last *k*-mer of *u_c_* and first(*v*) corresponding to the first *k*-mer of *v*. We define the edge weight as 1 – *J* (last(*u_c_*), first(*v*)). We continue the traversal with the unseen successor that has the edge with the lowest weight

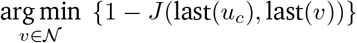

If *u_c_* has successors but all of them are seen, the DFS recursion steps back and continues along the next best edge. The traversal step is recursively repeated until it reaches a sink. In this case, we reset all traversal states in *G* to unseen and continue with the next source vertex. Intuitively, this traversal scheme favors paths in the graph that have a high color identity among consecutive vertices.

## 3 Results

We implemented the methods described in Section 2 in PopIns2 using the *Bifrost* [Holley and Melsted, 2020] and *SeqAn* [Reinert *et al.*, 2017] C++ libraries. The program comprises successively executable submodules for the assembly, merging, placing, and genotyping steps. We assess PopIns2 on simulated data and compare its results to PopIns and Pamir [Kavak *et al.*, 2017] as they are, to the best of our knowledge, the only other programs that are specifically tailored to identifying NRS sequences on many genomes simultaneously. We further demonstrate the feasibility of running PopIns2 on 150 samples from the Polaris Diversity Cohort (BioProject accession PRJEB20654) and assess the predicted NRS variant genotypes based on inheritance patterns using the Polaris Kids Cohort. Finally, we applied the new merge algorithm to whole-genome sequencing data of 1000 Icelanders.

### 3.1 Detecting NRS in simulated data

We implemented a workflow to simulate short-read sequencing data from human chromosome 21 (downloaded from ftp://ftp.ncbi.nlm.nih.gov/genomes/all/GCA_000001405.15_GRCh38/seqs_for_alignment_pipelines.ucsc_ids/GCA_000001405.15_GRCh38_no_alt_analysis_set.fna.gz on November 8th, 2016). In this workflow, a set *I_sim_* of 500 sequences (length mean=1455.17, sd=1526.08, min=41, max=9190, non-overlapping and in a distance of at least 1000 bp from N-sequence) is cut from chromosome 21 and the modified chromosome (*chr*21^−^) is further used for generating data of diploid individuals. The data is generated in two steps. First, two new haplotypes *h*_1_, *h*_2_ are created by insert-ing sequence subsets 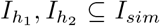 into two copies of *chr*21^−^. Second, short-reads are simulated [Huang *et al.*, 2012] from both *h*_1_, *h*_2_ and mapped [Li, 2013] to *chr*21^−^ as the reference. The sequences 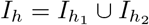 are NRS with respect to *chr*21^−^. This simulation workflow does not introduce artificially generated sequences but only real human sequences that originated from human chromosome 21.

We simulated data for 100 diploid individuals at 30x coverage using this workflow and applied PopIns, PopIns2 and Pamir to the data. All three tools yield a FASTA file containing the NRS found across all individuals (NRS sets). We compare each NRS set to the truthset 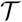, which is defined as

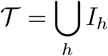

To assess a NRS set 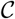 with respect to 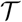 we performed an all-vs-all sequence alignment with minimum alignment length of 50 base pairs using STELLAR [Kehr *et al.*, 2011], i.e., we align every sequence in 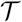 with every sequence in 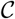. We use a bipartite matching procedure (Supplementary Figure 2) to compute precision and recall. Every 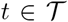 that is at least 90% covered by at least one individual alignment with a sequence 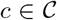 is counted as a true positive (TP). Every other 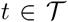 is counted as false negative (FN). The number of false positives (FP) is the difference of the set cardinality 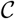 minus TP. This implies that, as opposed to less stringent prior analysis [Rizk *et al.*, 2014, Kehr *et al.*, 2016], we count redundant alignments as FP. The number of redundant alignments is calculated as the difference of all 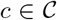 that cover 90% of a *t* with an alignment minus TP. The *F*_1_ score is the harmonic mean of precision and recall.

For every simulated sample we used the *PopIns2 assemble* module to create contigs from unmapped reads as it offers more flexibility than the previous assemble module in PopIns. It was used with Minia [Drezen *et al.*, 2014] as internal assembler (default) and with Velvet to mimic the *PopIns assemble* module.

#### Impact of the ASF parameter on the assembly step

We noticed that the ASF parameter of the assemble module plays a key role for the amount of reads being classified as unmapped and evaluated different settings of the ASF parameter in Supplementary Table 1. Setting a higher ASF value causes a less stringent selection of unmapped reads, i.e., fewer base pairs of a mapped read in relation to its read length have to be classified as unmapped for the read to be classified as unmapped. Interestingly, within the range of tested values, more unmapped reads resulted in fewer contigs per sample.

#### Evaluation of the merging step

We used the PopIns and PopIns2 merge modules with the different contig assemblies and Pamir to generate sets of NRS and evaluate their precision and recall with respect to 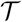. The NRS detected by Pamir (called *events*) are written with 1000 bp flanking regions of the genome. To not penalize the alignments for the flanking regions of the NRS we trimmed the flanking regions prior to the computation of recall and precision.

Supplementary Tables 2 and 3 summarize the results of the different merging approaches for 50 and 100 simulated samples. The combination of PopIns2 and contigs assembled with Minia3 shows the highest precision among all tested setups while Pamir has the highest recall. Velvet combined with the PopIns2 merge module was not tested extensively since early results indicated that Minia3 outperforms Velvet for our purpose of NRS assembly. We tested three different setups for the ASF parameter of the PopIns2 assemble module: *0 .5* (PopIns default), *0 .67* (PopIns2 default) and *0 .75* (very soft read filtering). With an increasing ASF and using 50 simulated samples, PopIns and PopIns2 showed a total gain in *F*_1_ scores of 16 percentage points and 18.7 percentage points, respectively. Among all three tested approaches the best performing setups show a difference in *F*_1_ scores of 5.5 (n=50 samples) and 4.8 (n=100 samples) percentage points.

We also investigated the NRS sets for different numbers of samples (Figure 1). We used a maximum of 100 samples and tested the default parameters of every NRS variant caller as well as PopIns2 with an increased ASF value. PopIns2 has the best recall for sample numbers up to 10-20. Then Pamir outperforms the best PopIns2 setup (by 2.2 to 4.7 percentage point). The recall value of PopIns2 is virtually constant at 0.68 to 0.73 depending on the setup. Pamir reveals a substantial increase in recall from 0.29 to 0.77 within the first 20 samples. The precision of PopIns2 is again relatively constant at 0.77 to 0.86 depending on the setup with the exception of a slight drop (about 4 percentage points) at around 20 samples. Moreover, PopIns2 shows the highest precision in all ranges over 40 samples. The precision of Pamir is the highest among all approaches until approximately 40 samples (precision at 0.87), then PopIns2 performs the best until the maximum tested amount of 100 samples (up to 6.6 percentage points over Pamir). From its peak with very few individuals the precision graph of Pamir shows a slow linear decline with increasing sample numbers. We attribute this decline in precision primarily to the rapid increase of redundant sequences in the callsets of Pamir (see Supplementary Table 3).

Furthermore, we separated the true positives of PopIns2 and Pamir by the length of the NRS (Supplementary Figure 3) to investigate whether there exists a detection bias towards certain sequence lengths. In this experiment, all tested approaches reveal a length distribution approximately proportional to the truthset. Pamir showed slightly more true positives in the length categories up to 1000 bp while PopIns2 showed a few more in the categories from 1000 bp to 5000 bp. Note that PopIns2 cannot generate NRS sequences smaller than the *k*-mer size *k*. Due to the graph simplifications of the CDBG, even NRS sequences smaller than 2*k* – 1 bp might not being detected. PopIns2 was executed with the default *k* = 63.

### 3.2 Detecting NRS in the Polaris Diversity Cohort

To assess the new merging algorithm on real population-scale sequencing data we applied PopIns2 and PopIns to the Polaris Diversity Cohort (PDC). The PDC comprises the genomes of 150 individuals from three continental groups (AFR, EAS, EUR) short-read sequenced on an Illumina HiSeqX sequencer with a target whole genome coverage of 30X. We aligned the short-read data of every individual to the human reference genome (hg38) using BWA [Li, 2013]. We did not compare to Pamir on the PDC data as Pamir exceeded the maximum running time of 28 days on our compute cluster using 16 threads. Pamir takes the BAM files as input and runs the entire NRS variant calling at once.

#### Assembly step

The alignment files (BAM files) are used as input for the *PopIns2 assemble* module. PopIns2 assemble with default parameters produces an average of 8049 contigs per genome from an average of 7194710 unaligned or poorly aligned reads. For a fair comparison of the merging algorithms we used the same contig assemblies for use with PopIns and PopIns2. The contig assembly reduced the average disc memory requirement per individual from 1595 megabytes for the selected reads to six megabytes for the contigs.

Individual instances of *PopIns2 assemble* can be distributed across a high-performance compute cluster and support multi-threading of the most computation-intensive tasks (alignments and assembly). The 150 PDC instances of PopIns2 assemble were distributed across 16 compute nodes supporting the same CPU instruction set (Intel(R) Xeon(R) Gold 6130 CPU @ 2.10GHz) and took approximately 80 minutes of wall clock time per instance using 16 CPU threads.

#### Merging step

On the contigs of the PDC, PopIns2 merge generated 15306 NRS in 50 minutes of CPU time using a single thread. A break down of the running time of merging can be found in Supplementary Table 6. Building the CDBG and traversing the vertices are the dominating factors for the running time. By comparison, PopIns merge generated 13456 NRS in 94 minutes of CPU time using a single thread. The right panel of Figure 2 shows a monotonic increase in the number of NRS detected by each tool for growing numbers of genomes.

Next, we examined the scalability of the PopIns and PopIns2 merge algorithms by comparing required computing resources for growing numbers of genomes (Figure 2). For PopIns merge we observed a fast increase in memory consumption with increasing number of genomes. Merging the contigs for 150 genomes of the PDC required almost 100 gigabytes of main memory. In contrast, PopIns2 merge used a maximum of 342 megabytes of main memory for the same genomes and the CDBG fits into a 154 megabytes GFA file when written to disc. Both merge algorithms show a running time that is increasing linearly with the number of genomes. However, the running time of the merge algorithm of PopIns2 is increasing substantially slower than that of PopIns. Supplementary Figure 6 illustrates how the resource requirements of PopIns2 merge depend on its parameters. We chose the set of default parameters (k=63, g=49) for a fast running time and reasonably low memory consumption.

### 3.3 Genotype assessments using the Polaris Kids Cohort

In addition to the PDC, the Polaris project data includes whole-genome sequencing data for the Polaris Kids Cohort (PKC), a set of 50 individuals that are the *F*_1_ offspring of 100 parent pairs in the PDC. We use the 49 family trios (as of March 22nd, 2021, the sequence data of one kid could not be downloaded from the archive) to detect and genotype NRS variants and assess genotype concordance with Mendelian Inheritance rules and expected transmission rate.

#### Running PopIns and PopIns2 on the 49 trios

Analogously to the PDC, we first aligned the reads of 49 individuals in the PKC to the reference genome and used the resulting BAM files as input for *PopIns2 assemble*. PopIns2 assemble produced an average of 8187 contigs per individual.

We used the contigs from the previous step and the contigs of the 98 related individuals from the PDC for the *PopIns* and *PopIns2 merge* module. PopIns and PopIns2 generated 12889 and 15450 NRS, respectively, processing all 147 samples at once. In Supplementary Figure 7 and Supplementary Note 4 we explain an approach to compute set overlaps of the two joint NRS callsets of PopIns and PopIns2. The NRS sets overlap by 19-65% depending on the setup.

We also applied the *place* and *genotype* modules to the NRS of PopIns and PopIns2 and wrote the genotype predictions of all 147 individuals into one multi-sample VCF file using VCFtools [Danecek *et al.*, 2011].

#### Running Pamir on the 49 trios

As the trio data comprises the same number of genomes as the PDC, we did not expect Pamir to finish the NRS variant calling within the maximum run time on our compute cluster. Therefore, we ran Pamir separately with the genomes of one family trio at a time, i.e. we started 49 runs with three genomes each.

#### Variant counts and principal component analysis

We observed a median of 2256, 2463 and 1873.5 NRS variants per sample for PopIns, PopIns2 and Pamir, respectively. In Supplementary Figure 8 we provide NRS variant counts separated by the continental groups. Consistent with previous studies [Wong *et al.*, 2020, Abel *et al.*, 2020], the African (AFR) genomes exhibit the highest average number of NRS variants. Since we could not compute a joined callset of 147 individuals with Pamir we computed callset intersections for each trio separately (Supplementary Figure 9). Among the three pairwise intersections of NRS callsets PopIns and PopIns2 show the largest intersection.

In addition, we performed a principal component analysis (Supplement Figure 12) as a sanity check that our variant calls can be used to clearly distinguish samples from different continental groups. Analogously to previous call set evaluations [Niehus *et al.*, 2021], we converted the genotypes into a variant-sample matrix containing NRS variant allele counts and filtered uninformative NRS variants and those in linkage disequilibrium. As a result, the principal component analysis was calculated on 1787 variants. The first and second principal component successfully cluster the continental groups with 5.138% and 2.217% explained variance, respectively.

#### Mendelian inheritance error rate and transmission rate

We further examined the genotype predictions and the inheritance patterns of NRS variants in the 49 trios by utilizing the pedigree information. We did not include Pamir’s results into this analysis as Pamir does not report genotype qualities or likelihoods for the predicted genotypes.

For both PopIns and PopIns2 we calculated the Mendelian inheritance error rate and transmission rate as in [Niehus *et al.*, 2021]. The Mendelian inheritance error rate is a measure to assess the plausibility of variant genotypes in related individuals. It is the fraction of offspring genotypes that cannot be explained using Mendelian inheritance rules from the parental genotypes. The transmission rate is used as a measure to assess how often a variant allele is transmitted from parent to child. In the diploid human genome we expect a heterozygous carrier of a variant to transmit the variant by chance in 50% of the cases. We examined the transmission rate only in the cases where one parent was genotyped as heterozygous carrier and the second parent as a non-carrier. As heterozygous variants with low quality scores are particularly overabundant in both data sets, we additionally created variant subsets without variants that are not in Hardy-Weinberg Equilibrium (HWE) in the original callset.

Our analysis of Mendelian inheritance error rate and transmission rate (Supplementary Figure 10) shows that the NRS variant callsets of both PopIns and PopIns2 can be filtered to a Mendelian inheritance error rate below 1%. The amount of NRS variants per trio consistent with Mendelian inheritance rules is virtually identical for both methods under both HWE filter conditions. Under the condition of HWE filtered callsets PopIns shows a marginally better (≤0.25 percentage points) Mendelian inheritance error rate than PopIns2 at equal genotype quality thresholds. Under the same conditions PopIns2 maintains a marginally lower absolute deviation from the targeted 50% transmission rate than PopIns at equal genotype quality thresholds. The Mendelian inheritance error rates measured for Pamir have a median value of over 8.5%. We explain the data generated for Pamir in Supplementary Note 6.

### 3.4 Detecting NRS in 1000 Icelandic genomes

To further test the scalability of the new merging algorithm with a previously prohibitive number of genomes [Kehr *et al.*, 2017] we applied the PopIns2 merge module to a set of 1000 Icelandic WGS samples [Gudbjartsson *et al.*, 2015, Jónsson *et al.*, 2017].

The sets of unaligned and poorly aligned reads of each sample were previously generated with PopIns assemble. For each sample we reused these sets of reads and manually finalized the assembly step using Minia with the internal setup of PopIns2 assemble. The final assemblies contain 6301 contigs on average per genome.

The Minia contigs were input to the PopIns2 merge module and generated 61,515 NRS. Supplementary Figure 5 shows the length distribution of the NRS. The merging took 4 hours and 45 minutes wall clock time (3 days, 18 hours and 1 minute CPU Time) using 24 CPU cores and used a maximum of 2.47 gigabytes of main memory during the computation. A break down of the wall clock time of merging can be found in Supplementary Table 6.

## 4 Discussion

PopIns2 implements a scalable approach for generating a NRS variant call set from population-scale short-read whole-genome sequence data. We demonstrate on simulated data that the accuracy of PopIns2 meets that of previous tools while our new merging approach scales to orders of magnitude more input data.

The definition of poorly aligned reads is crucial for the assembly of NRS. We show that by raising the ASF in PopIns2 we can increase both recall and precision of NRS assembly on simulated data. However, we noticed in real data that a high ASF value leads to the selection of a large number of reads that may distort NRS assembly. Therefore, we chose a moderate ASF value of 0.67 as default value.

Our new approach for merging NRS from many genomes allows simultaneous processing of many genomes together. The approach is based on a CDBG and heavily relies on the color information. Processing only few genomes together (few colors) may lead to arbitrary traversal decisions while processing many genomes together leads to greater graph complexity. Still, we observe that the accuracy is robust across the tested range of sample numbers on simulated data suggesting that color-based traversal decisions counteract graph complexity.

The new approach leaves room for future extensions. For example, the paths through the graph have weights that may be used to compute a confidence score for each NRS. Furthermore, the traversal of the graph is trivially parallelizable on connected components of the graph. Analogously to other assembly approaches [Turner *et al.*, 2018], read pairs could be used to annotate long-range information in the graph. Finally, the greedy heuristic can be supplemented with further traversal rules and requirements for the sequences eventually output.

With the new merging approach, we addressed the scalability to large numbers of genomes. Future research may address other aspects of NRS variant detection, for example the accuracy of genotyping. In our evaluation on the Polaris data, two thirds of the variants were discarded by the HWE filter indicating an overabundance of heterozygous genotype predictions. We previously addressed this by introducing an alternative genotyping scheme [Kehr *et al.*, 2017] that assumes the NRS to be flanked by repeats. Incorporating both cases into one genotyping framework is a promising idea to obtain better genotype predictions for NRS variants. Also, recently it was shown [Chen *et al.*, 2019, Eggertsson *et al.*, 2019] that sequence graphs have tremendous potential to improve the genotyping accuracy for structural variants. We investigate those data structures for future versions of PopIns.

In the assessment on simulated data we used more stringent evaluation criteria than in previous studies in order to reveal differences between the tools on comparably simple simulated data sets. An alternative possibility for future work is to simulate data with more challenging NRS variants, e.g. NRS variants flanked by duplicated or inverted sequence as frequently observed in real data or NRS variants previously detected in real data. Further evaluations could also include the effect of the allele frequency and sequencing depth on NRS detection.

Short read data is still the most prevalent type of sequencing data. There is no doubt that long read and linked read data can resolve NRS variants even better [Meleshko *et al.*, 2019, Ebert *et al.*, 2021]. Combining the data from many genomes sequenced with the long read or linked read technologies will be a promising future research direction as more and more data sets become available [Beyter *et al.*, 2021].

While the largest number of sequenced genomes are human genomes and all our tests were performed on human data, our methods are not human-specific. We encourage application of PopIns2 to data sets of genomes from another species, in particular from other animals or plants.

## Acknowledgements

The authors would like to thank Brian Caffrey and Kedi Cao for their feedback on installing and running the software. Computation has been performed on the HPC for Research cluster of the Berlin Institute of Health.

## Funding

The project was supported by the Federal Ministry of Education and Research (BMBF) through grant number FKZ 031L0180 to B.K.

## Competing interests

G.H. and B.V.H. are employees of deCODE genetics/Amgen,Inc. W.T.J.W. is an employee of Google,Inc. The other authors declare no competing interest.

---

**Supplementary Figure 1.**
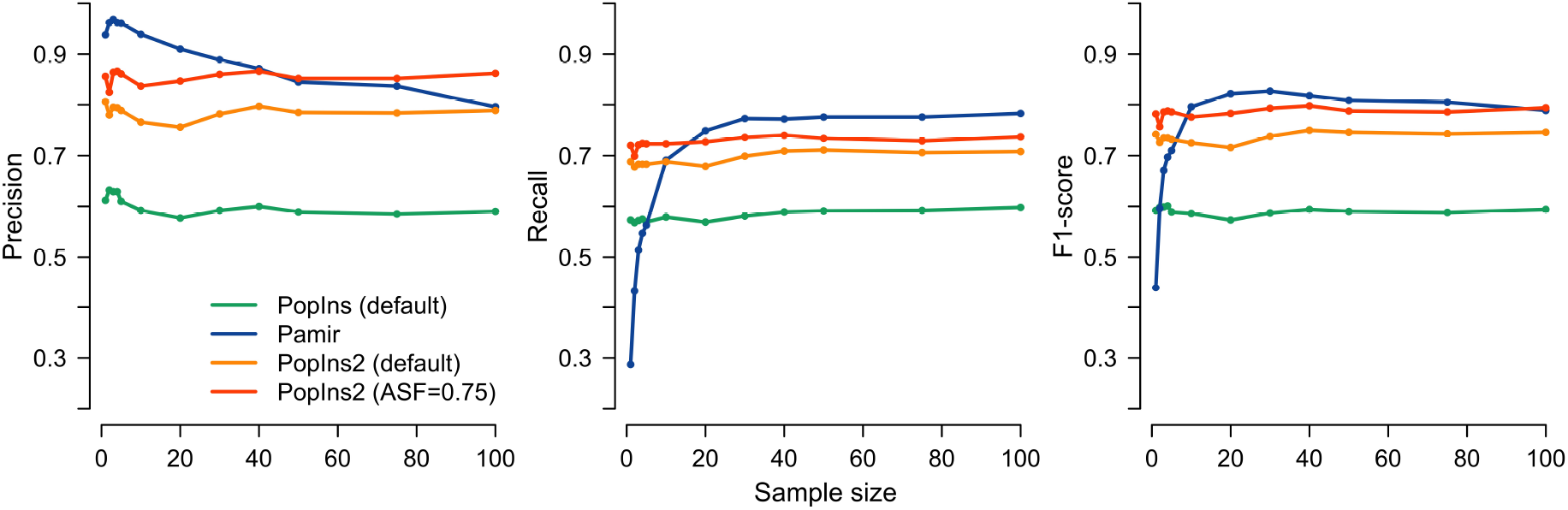
Precision, recall and F_1_ score of NRS callers on simulated data. NRS were assembled using a range of sample numbers (n = 1, 2, 3, 4, 5, 10, 20, 30, 40, 50, 75, 100).

**Supplementary Figure 2.**
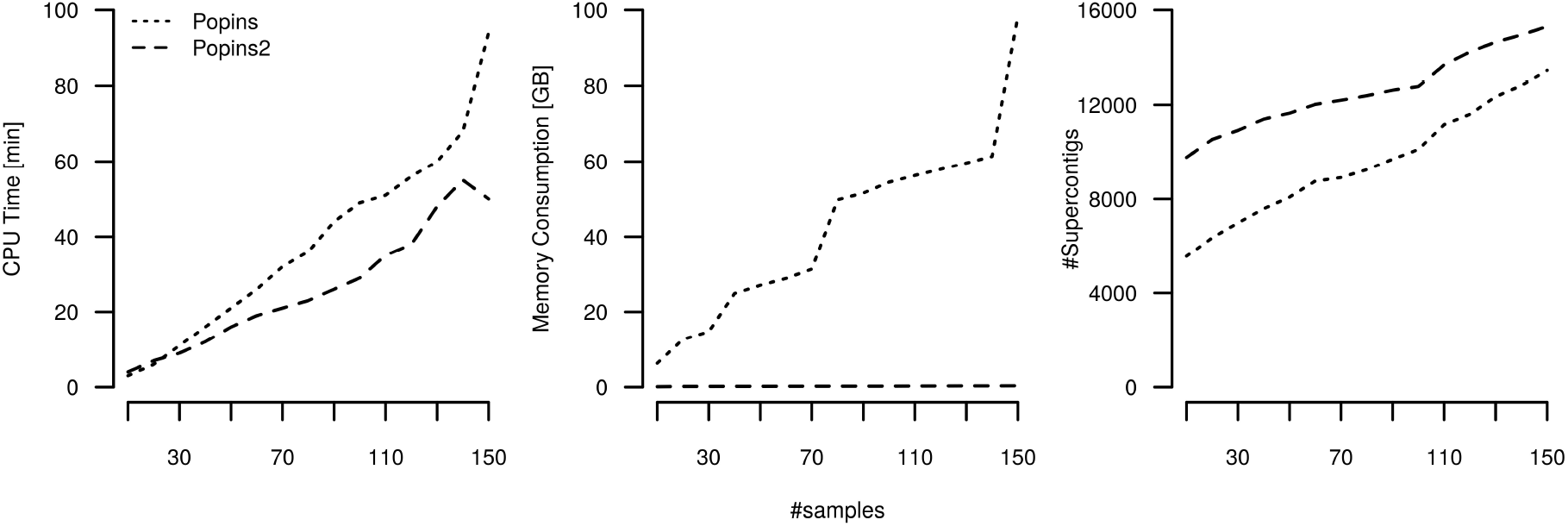
Benchmarks of the PopIns and PopIns2 merging modules on a growing subset of individuals from the Polaris Diversity Cohort. The left panel shows the CPU Time in minutes, the middle panel shows the memory consumption in gigabytes and the right panel shows the number of NRS.

## Notes

### Summary of Updates

Updated software New wording More statistics on the CDBGs (Supplements) Evaluation of KRAKEN2 to replace some internal steps of PopIns2 QUAST assembly assessment (Supplements)

