## Supplemental material for "Population-scale detection of non-reference sequence variants using colored de Bruijn Graphs"

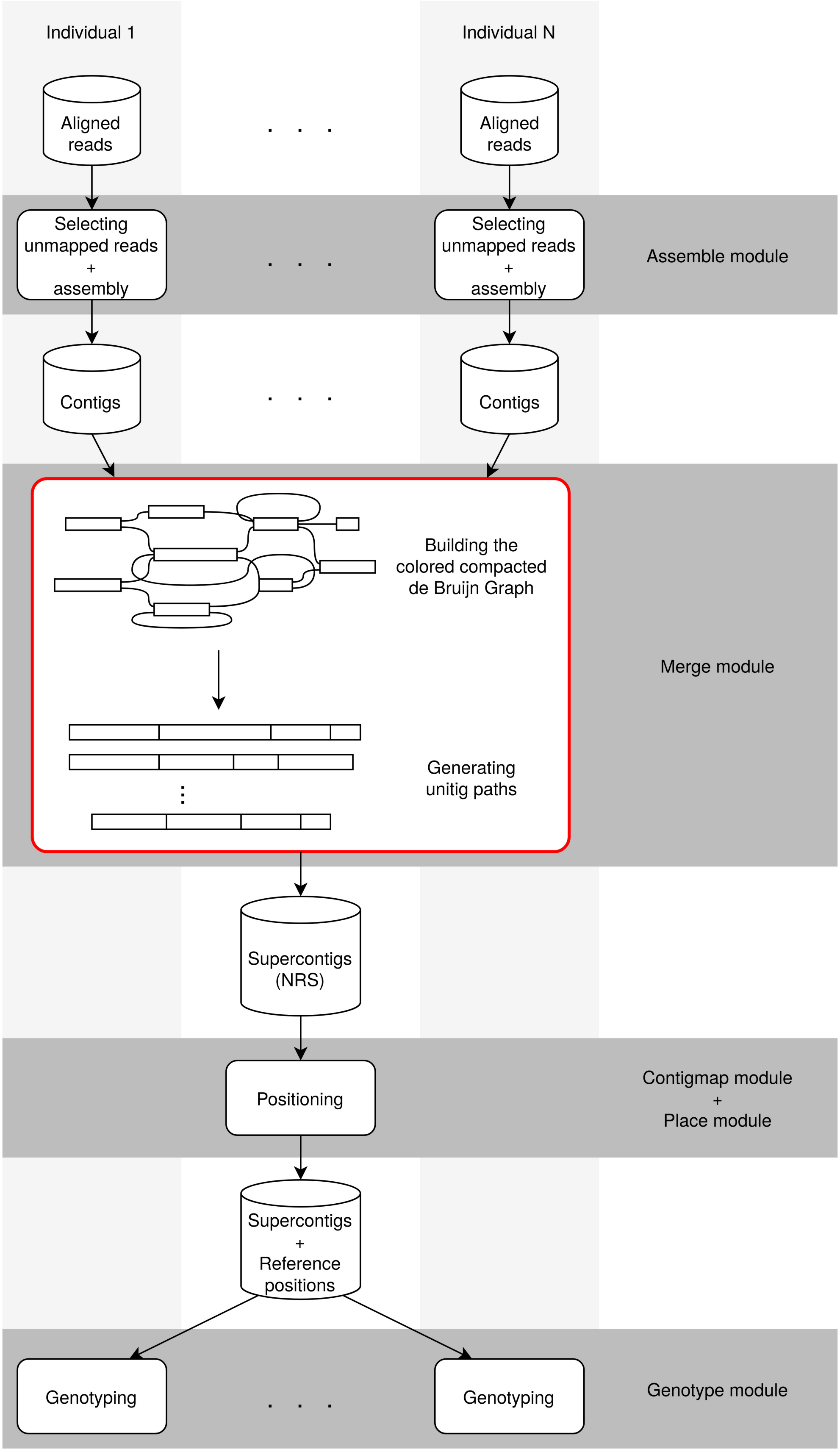


**Supplementary Figure 1.** *Overview of the PopIns2 workflow and its submodules. The merge module (denoted in red) was entirely revised compared to its predecessor PopIns.*

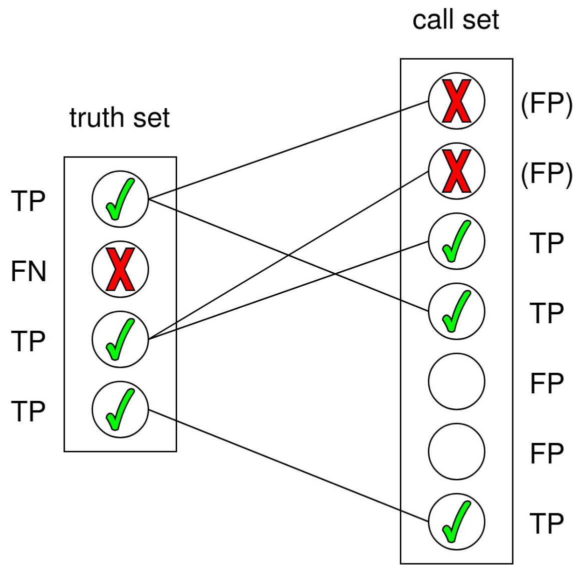


**Supplementary Figure 2.** *Bipartite matching graph to determine true positive (TP) and false negative (FN) findings with respect to the truthset and true positive, false positive (FP) and redundant ("(FP)") results from a callset from simulated data.*

**Algorithm 1** (Greedy Depth First Search in a dBG). *Given a colored de Bruijn Graph $G = (V, E, C)$, the Depth First Search (DFS) algorithm searches for the path with the highest consistent color annotation of every startnode $s \in V$ in G.*

*Let $u.id$ be a unique identifier of a unitig $u \in V$, $G.Adj[u]$ the list of all accessible vertexes of $u$ in $G$ and $u.color$ the color property of a node u. Further, every node maintains a discovery state $u.seen$ determining whether it has been visited by the DFS or not. The set $S$ is a global storage for unitig identifiers. W.l.o.g. in the Pseudo-code we assume only one consistent traversal direction, mind that in the real implementation we have to determine predecessors and successors with respect to the unitig's orientation as stored.*

```
DFS−Init(G)
    for each u ∈ G.V
        p <− empty path
        if isStartnode(u)
            if not DFS−Visit(u, p)
                p.append(u)
            for each u ∈ G.V
                u.seen = false
            if hasEnoughNovelKmers(p, S)
                for each q ∈ p
                    S.add(q)
                print(p)
        if isSingleton(u)
            p.append(u)
            if hasEnoughNovelKmers(p, S)
                S.add(u)
                print(p)

DFS−Visit(u, p)
    if u.seen == false
        u.seen = true
        if G.Adj[u] is empty:
            p.append(u)
            return false
        G.Adj[u] <− sortByColors(G.Adj[u])
        for each v ∈ G.Adj[u]
            if not DFS−Visit(v, p)
                p.append(u)
                return false
    return true
```

| ASF | Avg. selected reads per sample | Avg. contigs per sample |
|---|---|---|
| 0.50 | 19647.20 | 152.2 |
| 0.67 | 20921.86 | 145.1 |
| 0.75 | 21358.84 | 140.2 |

**Supplementary Table 1.** *Average amount of selected reads and contigs per simulated individual resulting from the PopIns2 assemble module.*

### 1 Simulation and read alignment of the 100 simulated human samples

For each sample we used *ART_illumina* [Huang *et al.*, 2012] to simulate paired short-reads (Illumina HiSeq 2500, 150bp length, 15X read coverage per haplotype, 300bp fragment size with a standard deviation $\sigma$=50bp) from the simulated haplotypes of chromosome 21. The simulated reads in the resulting FASTQ files were aligned to the modified human genome (HGCh38 with missing sequences $I_{sim}$, see section 3.1) with *bwa mem* [Li, 2013]. Therefore, our simulation workflow considers alignment artifacts, i.e., some reads are not aligned to their original location on chromosome 21 but elsewhere in the genome. We call these reads *off-target* reads. On average each sample contains 7.49 million reads, 7.38 million of them are aligned or partially aligned to chromosome 21 and 12.7 thousand reads are labelled as unmapped. The remaining average number of 99.1 thousand reads are off-target reads. Derived from Supplementary Table 1, the ASF parameter settings 0.50, 0.67 and 0.75 lead to an average of 35.36%, 41.84% and 44.07% of the reads selected for assembly having a (poor) alignment to the reference genome, respectively.

| Caller | Assembler | #Individuals | ASF | Recall | Precision | $F_1$ score |
|---|---|---|---|---|---|---|
| PopIns | velvet | 50 | 0.50 | 0.623 | 0.536 | 0.576 |
| PopIns | minia3 | 50 | 0.50 | 0.591 | 0.589 | 0.590 |
| PopIns | minia3 | 50 | 0.67 | 0.715 | 0.748 | 0.731 |
| PopIns | minia3 | 50 | 0.75 | 0.742 | 0.758 | 0.75 |
| PopIns2 | minia3 | 50 | 0.50 | 0.589 | 0.616 | 0.602 |
| PopIns2 | minia3 | 50 | 0.67 | 0.709 | 0.781 | 0.743 |
| PopIns2 | minia3 | 50 | 0.75 | 0.734 | **0.854** | 0.789 |
| Pamir | - | 50 | - | **0.776** | 0.837 | **0.805** |
| PopIns2 | minia3 | 100 | 0.67 | 0.708 | 0.789 | 0.746 |
| PopIns2 | minia3 | 100 | 0.75 | 0.737 | **0.862** | **0.794** |
| Pamir | - | 100 | - | **0.784** | 0.747 | 0.765 |

**Supplementary Table 2.** *Precision and recall of NRS detection from simulated human chromosome 21 data of 50 and 100 diploid individuals.*

| Assembler | SV caller | samples | ASF | #elements covered from truthset | $\lvert$Callset$\rvert$ | #elements being aligned from callset | #elements being redundant from callset |
|---|---|---|---|---|---|---|---|
| velvet | popins | 50 | 0.50 | 297 | 554 | 298 | 1 |
| minia3 | popins | 50 | 0.50 | 282 | 479 | 286 | 4 |
| minia3 | popins | 50 | 0.67 | 341 | 456 | 347 | 6 |
| minia3 | popins | 50 | 0.75 | 354 | 467 | 370 | 16 |
| minia3 | PopIns2 | 50 | 0.50 | 281 | 456 | 281 | 0 |
| minia3 | PopIns2 | 50 | 0.67 | 339 | 432 | 341 | 2 |
| minia3 | PopIns2 | 50 | 0.75 | 350 | 411 | 355 | 5 |
| - | pamir | 50 | - | 370 | 442 | 418 | 48 |
| minia3 | PopIns2 | 100 | 0.67 | 347 | 440 | 349 | 2 |
| minia3 | PopIns2 | 100 | 0.75 | 361 | 419 | 365 | 4 |
| - | pamir | 100 | - | 384 | 514 | 480 | 96 |

**Supplementary Table 3.** *Counts of the precision and recall statistics of Supplementary Table 2. The truthset of the simulation contains 477 contigs at 50 samples and 490 contigs at 100 samples. An element t of the truthset is accounted as covered ($5^{th}$ column) if there exists at least one supercontig that aligns to at least 90% of t. In reverse, a supercontig is accounted as aligned if it can contribute an alignment that covers least 90% of a t ($7^{th}$ column). Mind that the amounts of covered t and aligned supercontigs are not necessarily equal since the callsets can have redundant sequences. The operator* $|\cdot|$ *denotes the set cardinality ($6^{th}$ column).*

| Variant caller | ASF | N50 | NGA50 | misassemblies | misassembled contigs | local misassemblies | mismatches per 100kbp | Largest alignment | Genome fraction [%] |
|---|---|---|---|---|---|---|---|---|---|
| PopIns | 0.50 | 1742 | 1403 | 17 | 16 | 0 | 30.92 | 7689 | 78.165 |
| PopIns2 | 0.50 | 1671 | 1200 | 13 | 12 | 1 | 6.49 | 7689 | 77.694 |
| PopIns2 | 0.67 | 1937 | 1500 | 23 | 23 | 1 | 11.01 | 9074 | 81.118 |
| PopIns2 | 0.75 | 2118 | 1700 | 33 | 26 | 6 | 7.20 | 8825 | 82.171 |
| Pamir * | - | 1869 | 1217 | 0 | 0 | 0 | 17.39 | 9073 | 71.991 |
| Pamir | - | 2980 | 1408 | 1 | 1 | 113 | 234.54 | 9074 | 73.029 |

**Supplementary Table 4.** *Assembly metrics computed via QUAST for 100 simulated samples and contigs of minimum 125bp length. The truthset $\mathcal{T}$ (main text section 3.1) was used as reference genome for QUAST. The assessment marked with "*" denotes the callset of Pamir where the flanking reference sequences were manually trimmed from the non-reference sequences. The legacy version of PopIns uses Velvet as internal assembler.*

### 2 Assembly metrics using QUAST

Supplementary Table 4 supports the simulated data analysis in section 3.1 of the main text. We used QUAST [Gurevich *et al.*, 2013], a quality assessment tool for genome assemblies, to assess the consensus of the supercontigs with the truthset $\mathcal{T}$.

The results in Supplementary Table 4 show that the chosen default value for the ASF parameter (0.67) is a reasonable middle ground between assembly continuity, number of misassemblies and covered sequence fraction by the callset. For PopIns2, the N50, the NGA50 but also the number of misassemblies grows with an increasing ASF. The continuity and accordance with the truthset of PopIns' callset ranks between the results of PopIns2 with ASF=0.50 and ASF=0.67. However, directly compared to the results of PopIns2 with matching ASF=0.50, the supercontigs of PopIns contain more misassemblies, misassembled contigs and mismatches. PopIns2 has less mismatches in its assemblies and covers a higher fraction of the truthset than Pamir with all tested ASF settings. However, Pamir seems to create less misassemblies. Note that the number of misassemblies in this analysis is not trivial to compare as all callsets (except for the manually cropped one; marked with "*") have flanking reference sequences. The comparison of the two callsets of Pamir shows that QUAST reports many more misassemblies if flanking regions are present.

Interestingly, both variant caller report a largest alignment of 9074 bp which is the length of the second largest sequence in $\mathcal{T}$.

| Cohort | #samples | ASF | #nodes | #edges |
|---|---|---|---|---|
| simulated | 50 | 0.50 | 500 | 117 |
| simulated | 50 | 0.67 | 478 | 127 |
| simulated | 50 | 0.75 | 476 | 183 |
| simulated | 100 | 0.50 | 527 | 165 |
| simulated | 100 | 0.67 | 521 | 231 |
| simulated | 100 | 0.75 | 537 | 327 |
| Polaris Diversity Cohort | 150 | 0.67 | 797236 | 2,195,847 |
| Polaris Trios | 150 | 0.67 | 817052 | 2,255,221 |

**Supplementary Table 5.** *De Bruijn Graphs created by the PopIns2 merge module.*

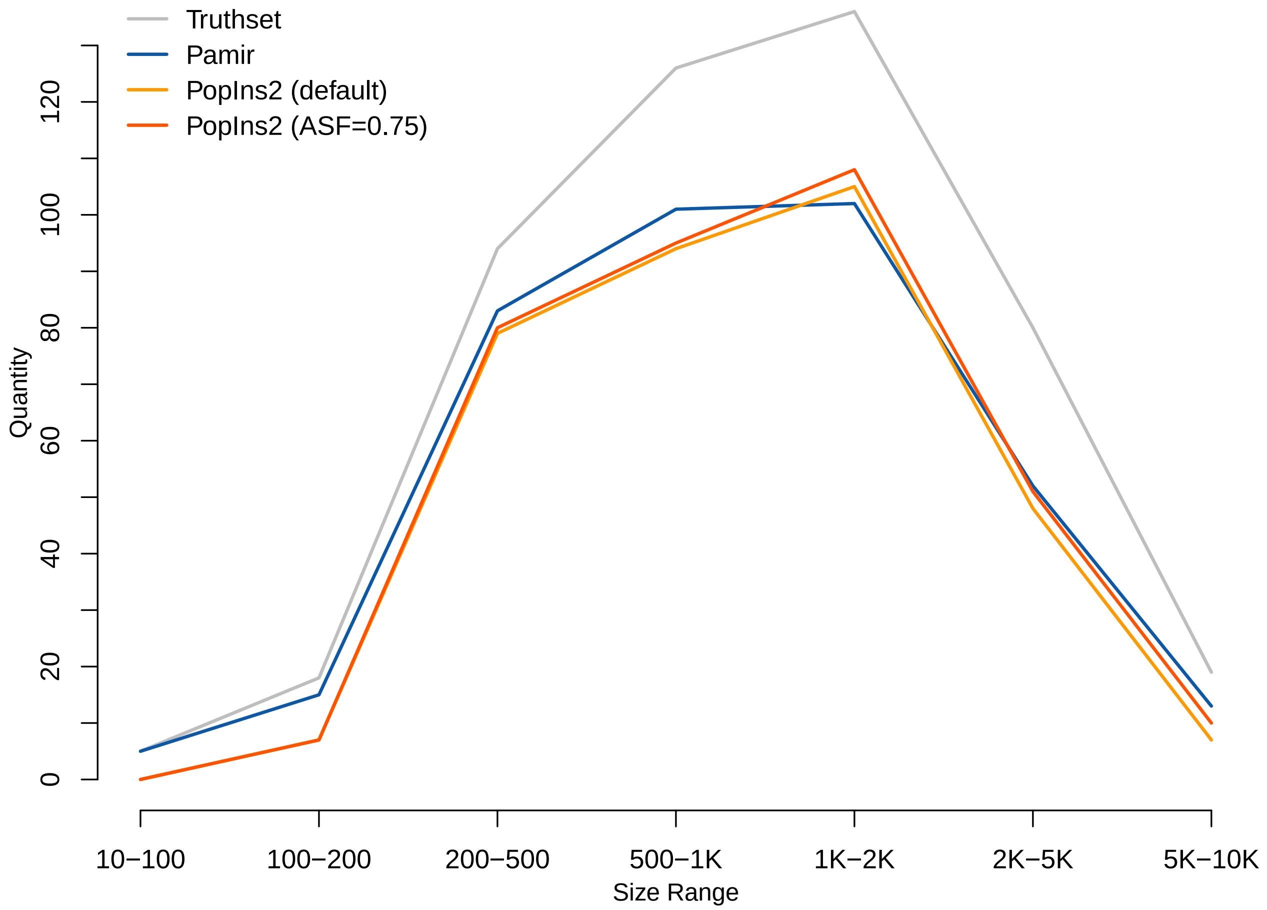


**Supplementary Figure 3.** *True positive counts for different size ranges of insertion sequences. The graph shows the counts of the ground truth (grey), PopIns2 and Pamir in their default setup and additionally PopIns2 with increased ASF. All counts were generated using 50 simulated samples.*

| Function | Wall clock time [min] | |
|---|---|---|
| | Polaris Diversity Cohort | Icelandic Genomes |
| Graph build (threads) | 35 (1) | 3 (24) |
| Graph simplification | 3 | 1 |
| Graph color annotation | 2 | 1 |
| Graph traversal | 98 | 220 |
| Other | 1 | 1 |

**Supplementary Table 6.** *Wall clock times measured in minutes for PopIns2 merge.*

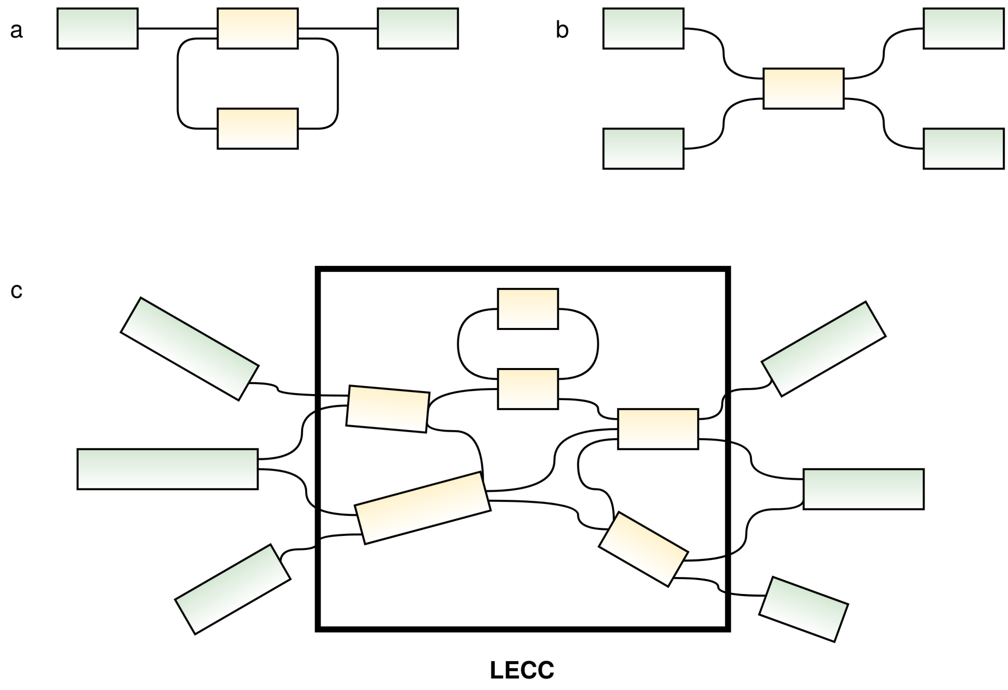


**Supplementary Figure 4.** *Topological sketches of a compacted de Bruijn Graph. Green and yellow nodes denote unitig sequences with an entropy higher or lower than a threshold x, respectively. Panel* ***(a)*** *shows a small low entropy repeat. Panel* ***(b)*** *shows a low entropy unitig with a bifurcation at both ends. Panel* ***(c)*** *shows a compacted dBG comprising both topological sub types (a) and (b). We call an interconnected group of low entropy unitigs a Low Entropy Connected Component (LECC).*

### 3 Bridging low entropy sequences

Many genomic sequences contain sub regions of low entropy (e.g. short tandem repeats, homopolymers, poly-A tails). Those sub regions have a prominently reduced variety in dinucleotides. Sampling k-mers from low entropy sequences typically results in many highly similar k-mers. Subsequently, when a dBG is built from highly similar k-mers they tend to form characteristic topologies (Supplementary Figure 4a,b). We observed that these topologies can grow and cluster into large, complex and densely interconnected components (Supplementary Figure 4c) which we term Low Entropy Connected Components (LECC). The traversal scheme of *PopIns2 merge* can make poor neighbor decisions if highly abundant k-mers across many samples form complex LECCs and potentially returns incorrect supercontigs, i.e. the supercontigs do not contain accurately assembled non-reference sequence. When *PopIns* was developed the same problem with low entropy sequences was discovered even though the merge algorithm is entirely different. The solution of PopIns to this problem is a threshold $y$ that leaves contigs below an entropy of $y$ ignored within the merge algorithm. For *PopIns2* we designed a mechanism that masks uncertainty in the supercontig sequences due to low entropy during the generation of a sequence.

**Finding partners around LECCs.** In the merge module prior to the traversal all unitigs with a sequence entropy below a user defined threshold (parameter *-e* with default=0, no consideration of LECCs) are annotated. Next, connected components of annotated unitigs (hence LECCs) are identified. For each LECC $L$ all unitigs $\mathcal{U}_L$ which border $L$ are stored together with the information which k-mer $k_L$ (head or tail) connects to $L$. Then, for every $u \in \mathcal{U}_L$ a DFS is initiated and searches through $L$ until all accessible unitigs $\mathcal{P} \subseteq \mathcal{U}_L$ (called "Partners") are found. The color vector of k-mer $k_L \in u$ is compared to the color vector of every $k_L \in p$, where $p \in \mathcal{P}$, using the Jaccard Index of k-mer colors. As a result, a map $\mathcal{M}_L$ stores the association between every $u$ and its best color matching partner $p$.

**Traversal with jumps.** Once the best color matching partners around the LECCs are computed the merge module continues with the main traversal method. If the traversal detects a LECC $L$ at some unitig $u$ during its recursive progression the algorithm makes a lookup in $\mathcal{M}_L$ to determine the partner of $u$. Then, instead of accessing a direct neighbor of $u$ the traversal jumps across $L$ directly to $p$. Later when the concatenation function $\omega$ generates the supercontig sequence from a given path, a jump over a LECC is encoded as a sequence of 'N' characters of length $k$.

**Adjustment of sequence masking.** The decision and extend of masking sequence to avoid potential unnoticed errors in supercontig sequences to left the user. By default PopIns2 does not apply masking and returns only complete sequences. However, insertions sequences that were generated from a path that led through a LECC tend to be less reliable in terms of sequence content and length. This becomes particularly severe if the true sequence contains STRs. As the merge module of PopIns2 is not optimized for accurate repeat resolution a false amount of copies of the repeat pattern can go unnoticed if the STR is longer than $k$ and low entropy is not masked. By comparison, the merge process of PopIns ignores all input contigs below an entropy threshold of 0.75 by default.
*PopIns2 merge* offers an option (command line parameter *-l*) to explore the masking process by writing a CSV file that contains the IDs of low entropy unitigs according to the entropy parameter *-e*. This CSV file can be loaded into Bandage [Wick *et al.*, 2015] to visualize the extend of LECCs in the graph.

**Complexity of finding partners around LECCs.** The mechanism to find partners around LECCs introduces additional computations to the merge algorithm but, in practise, its cost in run time is negligible compared to the main routine. The worst-case run time of our implementation to find all partners for all LECCs is

$$\begin{aligned} &\mathcal{O}(N(V + B(V_{LECC} + E_{LECC} + V + B \cdot C + B))) \\ =&\mathcal{O}(N \cdot V + N(B(V_{LECC} + E_{LECC} + V) + B^2 \cdot C + B^2)) \end{aligned}$$

where $V$ is the amount of vertices in the ccdBG, $E$ is the amount of edges in the ccdBG, $N$ is the amount of LECCs, $V_{LECC}$ is the amount of vertices in a LECC, $E_{LECC}$ is the amount of edges in a LECC, $B$ is the amount of vertices that border a particular LECC and $C$ is the amount of colors in the ccdBG. Supplementary Table 7 shows a low level running time analysis for the methods involved into the partner finding mechanism.

| Function | Run time |
| --- | --- |
| get_borders() | $\mathcal{O}(V)$ |
| DFS() | $\mathcal{O}(V_{LECC} + E_{LECC})$ |
| reset_def_states() | $\mathcal{O}(V)$ |
| check_accessibility() | $\mathcal{O}(V_{LECC} + E_{LECC} + V)$ |
| *calls DFS()* | $DFS()$ |
| *calls reset_def_states()* | $reset\_def\_states()$ |
| color_overlap() | $\mathcal{O}(C)$ |
| find_jumps() | $\mathcal{O}(N(V + B(V_{LECC} + E_{LECC} + V + B \cdot C + B)))$ |
| *iteration over all LECCs* | $N$ |
| *calls get_borders() per LECC* | $\mathcal{O}(N \cdot V)$ |
| *calls check_accessibility() per border* | $\mathcal{O}(B(V_{LECC} + E_{LECC} + V))$ |
| *calls color_overlap() for all border pairs* | $B \cdot B \cdot \mathcal{O}(C)$ |
| *reset partner accessibility per border* | $B \cdot B$ |

**Supplementary Table 7.** *Detailed listing of the contributions to the worst-case complexity of the LECC finding process.*

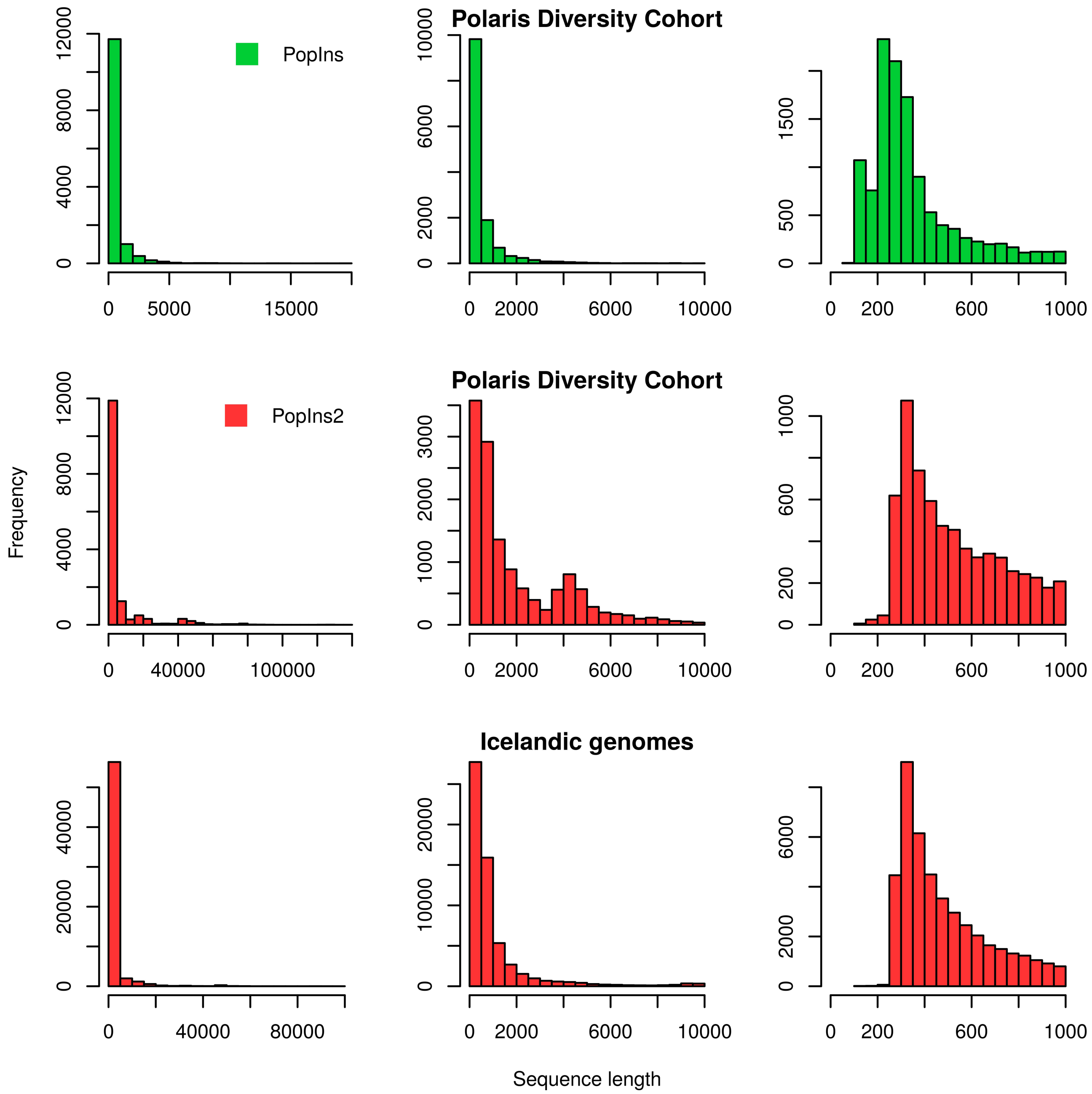


**Supplementary Figure 5.** *Lengths histograms of the supercontig in the Polaris Diversity Cohort and the set of 1000 Icelandic human genomes. The top (PopIns) and middle row (PopIns2) show the supercontigs of the Polaris Diversity Cohort and the bottom row (PopIns2) shows the set of Icelandic human genomes. Each row displays the same data three times (columns) restricted to different maximum lengths, i.e. all supercontigs are shown in the left column, supercontigs up to 10kbp are shown in the middle column and supercontigs up to 1kbp are shown in the right column.*

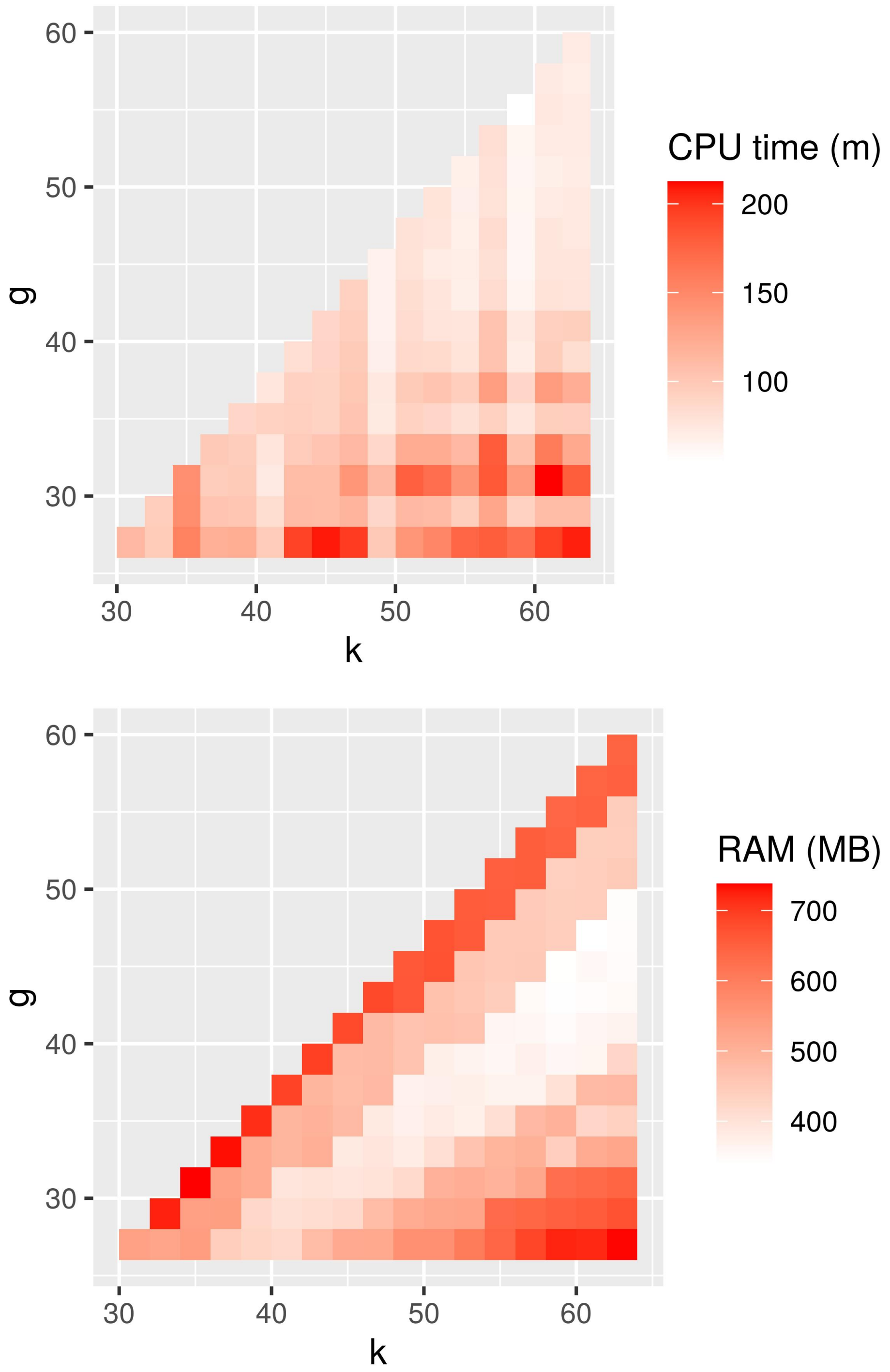


**Supplementary Figure 6.** *Parameter space exploration of kmer size (k) and minimizer size (g). Both parameters influence the construction of the colored de Bruijn Graph in PopIns2 merge. The heatmap at the top shows the CPU time measured in minutes for PopIns2 merge. The heatmap at the bottom shows the maximum memory consumption during the computation of PopIns merge in megabytes. Both heatmaps encode a high demand in computational resources in red and a low demand in computational resources in white.*

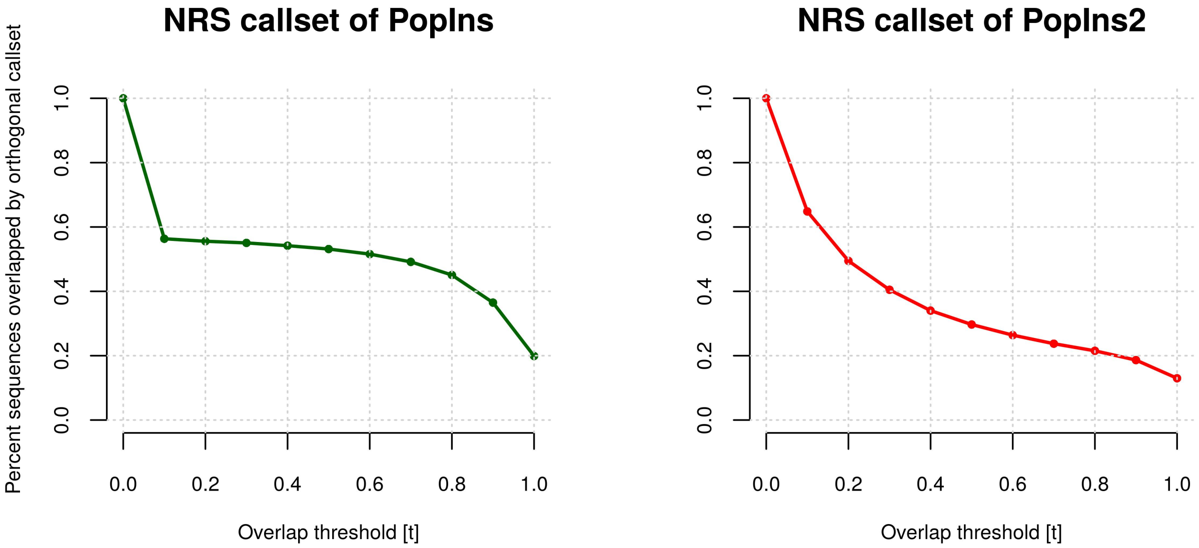


**Supplementary Figure 7.** *Set overlaps between the joint callsets of PopIns and PopIns2 from 147 samples of the PKC and its related PDC samples. The left panel shows how many sequences of the callset of PopIns are overlapped by a sequence of PopIns2 at minimum overlap threshold t. The right panel shows how many sequences of the callset of PopIns2 are overlapped by a sequence of PopIns.*

### 4 Set overlap of joint NRS callsets from 147 samples

Supplementary Figure 7 shows the diagrams for the set overlaps of the joint NRS callsets from PopIns (here set $P_1$) and PopIns (here set $P_2$). To determine a set overlap of $P_1$ and $P_2$ we first computed an all-vs-all sequence alignment (using STELLAR [Kehr *et al.*, 2011]; --*minLength 50*; maximal error rate --*epsilon 0.05*). With the given parameters we commonly observed a fragmentation of the pairwise alignments into multiple shorter local alignments. Hence, we decided to compute the set overlap separately for each callset as the amount of NRS that are covered by sequence of the orthogonal callset. In this context, a NRS $p_1 \in P_1$ ($p_2 \in P_2$) is covered if the fraction of base pairs that has no alignments to a $p_2 \in P_2$ ($p_1 \in P_1$) is below $1 - t$. We computed the set overlap of each callset for multiple minimum coverage thresholds *t*.

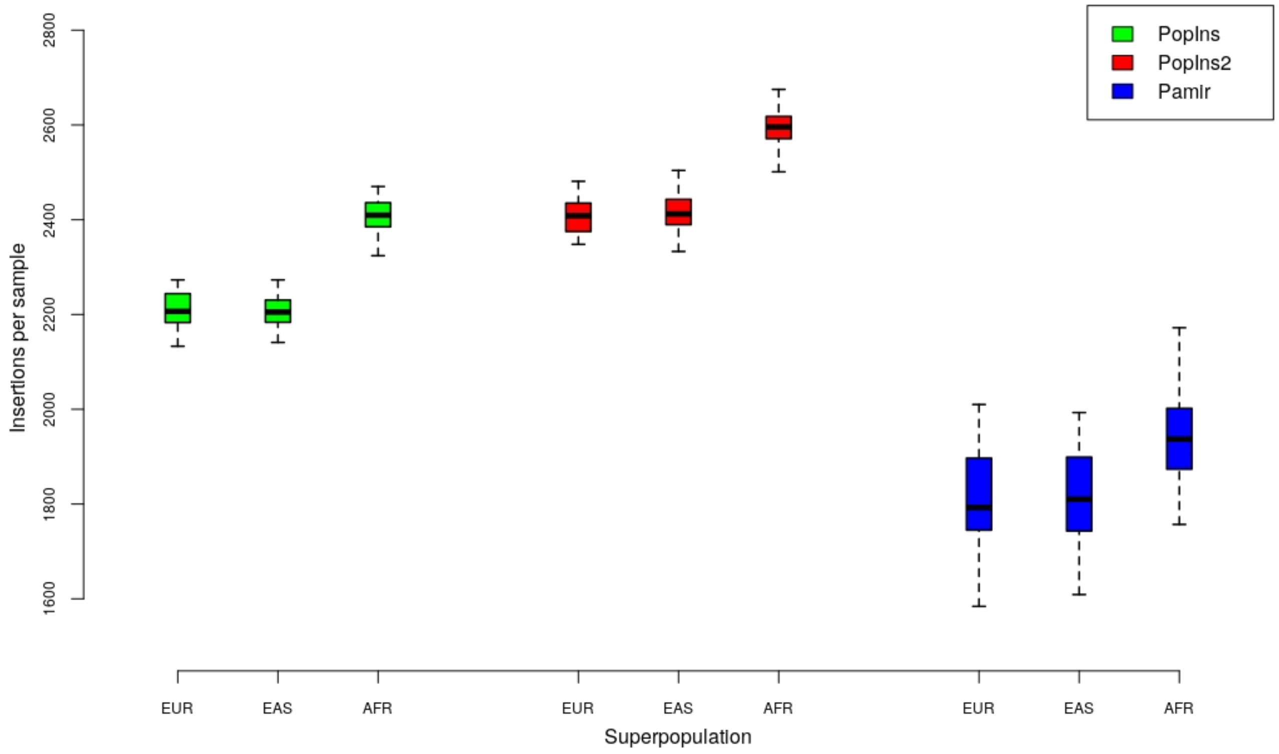


**Supplementary Figure 8.** *Number of insertions per sample and superpopulation of 147 related individuals from the PDC and PKC. PopIns and PopIns2 processed all 147 samples at once while Pamir processed three samples (one trio) at a time. The counts are taken from the final VCF files of PopIns, PopIns2 and Pamir and, consequently, each counted insertion was already placed and genotyped. Insertions are only counted if they are not genotyped with two reference alleles (0/0) and have a minimum length of 50 base pairs.*

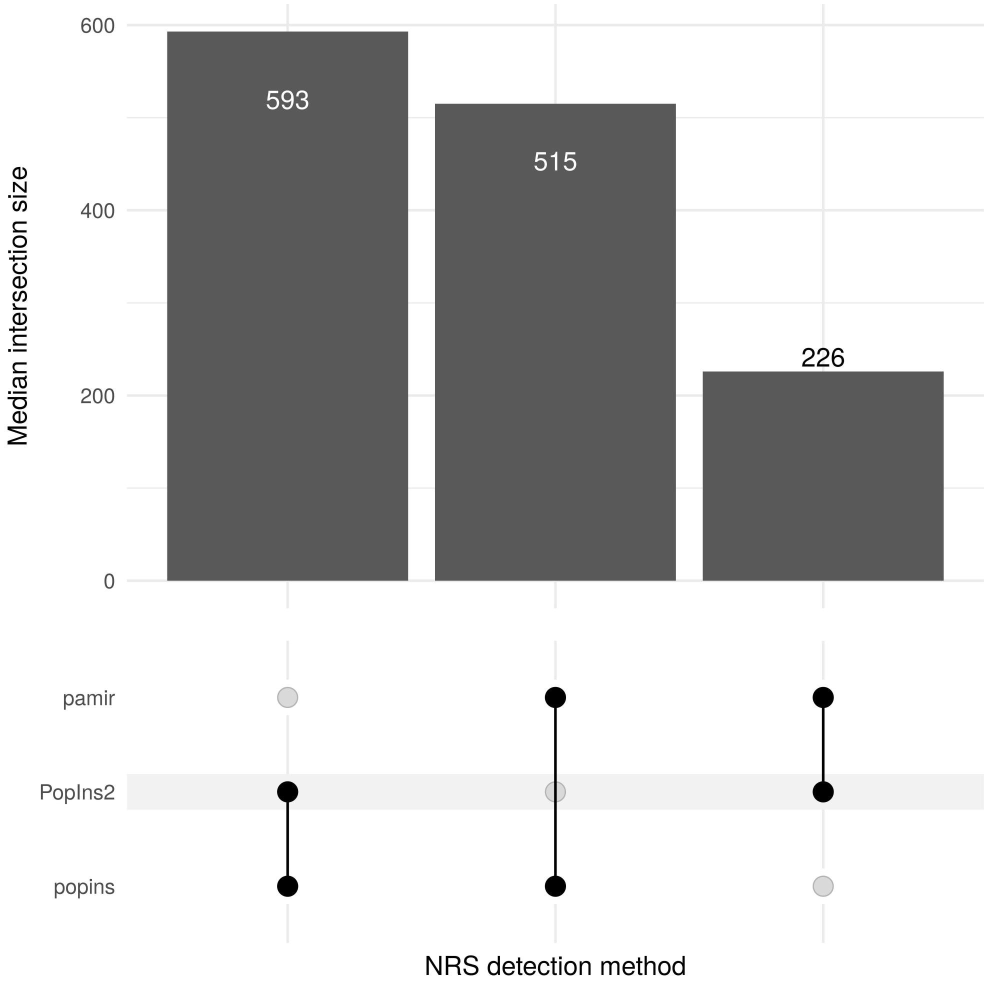


**Supplementary Figure 9.** *Set intersections of NRS callsets. The intersection sizes annotated in the plot are the median values of intersections computed for every trio pedigree and every pair of detection methods. The intersection of two NRS callsets of one particular trio is the number of distinct pairs of sequences with a reciprocal sequence overlap of at least 50% (using the same parameters as in Supplementary Note 4).*

### 5 Set overlap of NRS callsets from trio pedigrees

The NRS sequences in Supplementary Figure 9 were taken from the final VCF file of each caller and every trio pedigree. Hence, every NRS in this analysis was already (partially or fully) placed to the reference genome and genotyped. At least one individual of the trio pedigree needs to contain the variant in at least one allele for the NRS to be extracted from the VCF. Next, we trimmed the flanking reference sequences from the NRS of Pamir's callsets. We further excluded all remaining sequences of less than 50bp length from all callsets. The remaining callsets of Popins, PopIns2 and Pamir individually had a median value of 1650, 1898 and 6661 NRS sequences per trio, respectively. Finally, we used STELLAR [Kehr *et al.*, 2011] to compute the pairwise alignments.

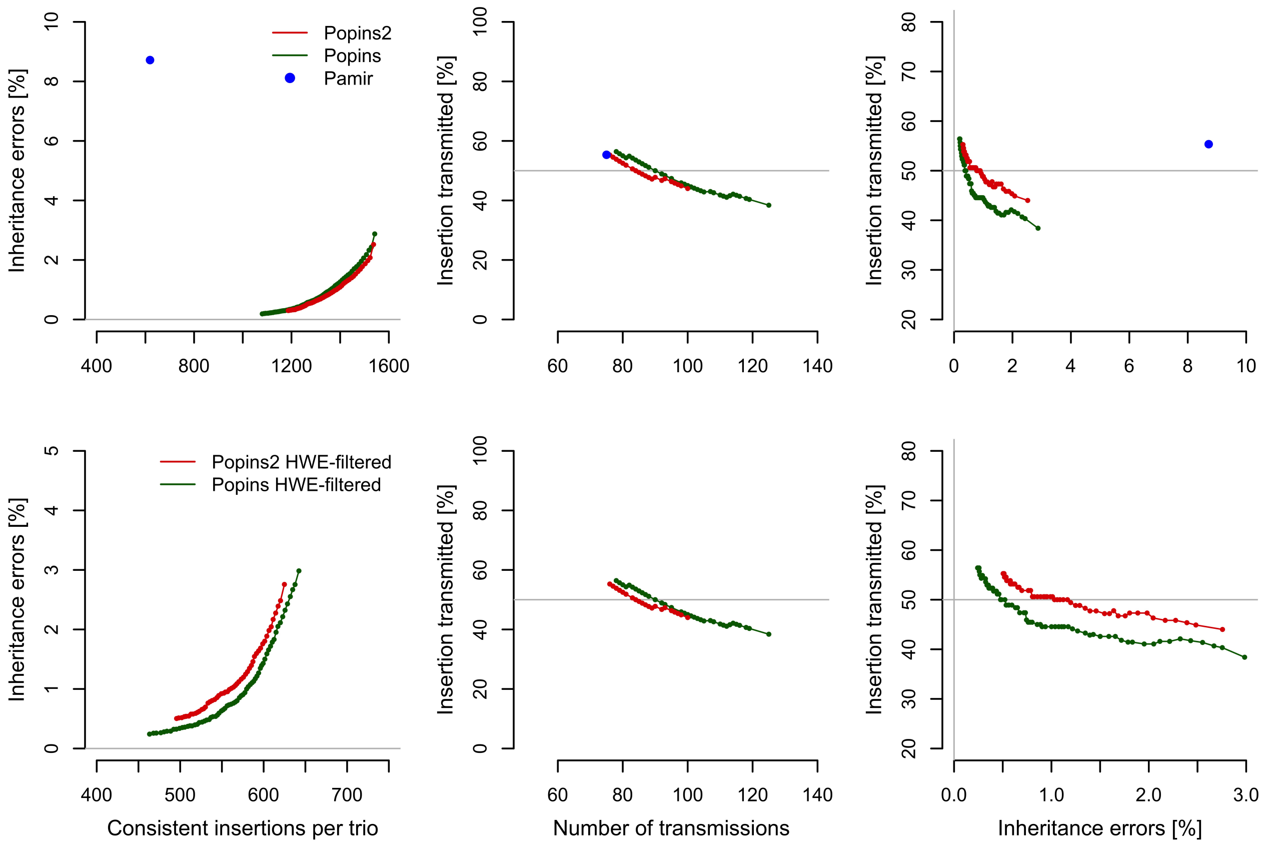


**Supplementary Figure 10.** *Mendelian inheritance patterns in the Polaris trios without (top row) and with filtering (bottom row) for Hardy-Weinberg-Equilibrium (P value 0.01). Mendelian inheritance error rate by amount of insertions per trio consistent with Mendelian inheritance rules (left panels). Transmission rate of insertions unique to one trio with one heterozygous parent by overall number of transmissions (middle panel). Transmission rate of insertions unique to one trio with one heterozygous parent by Mendelian inheritance error rate (right panel). The grey lines denote the ideal values. Each data point in the graphs corresponds to an observation of increasing minimum genotype quality.*

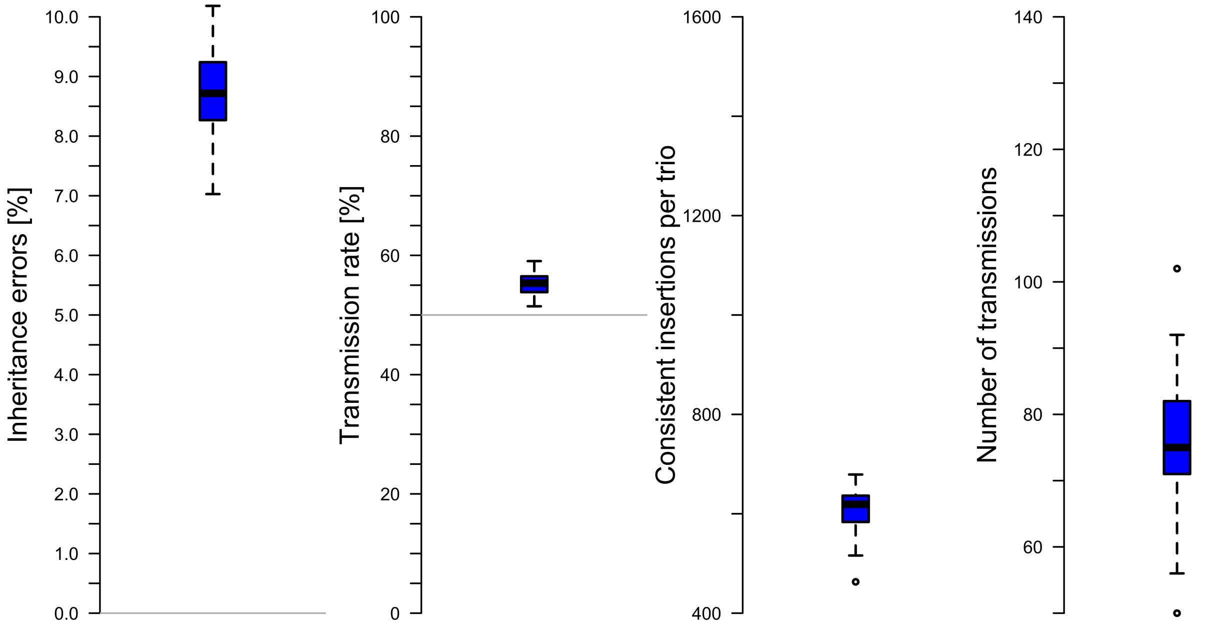


**Supplementary Figure 11.** *Distribution of observed Mendelian inheritance errors, transmission rate, consistent insertions per trio and number of transmissions from 49 trio pedigrees using Pamir. Grey lines denote ideal values.*

### 6 Mendelian inheritance error rate and transmission rate

In Supplementary Figure 10 we show the Mendelian inheritance error rate and transmission rate for different minimum genotype qualities (GQ) in the range [0,255]. Since Pamir only reports genotype predictions but no corresponding likelihoods or genotype qualities we only add one data point for Pamir into each plot. These data points for Pamir are the median values over the individual measurements of the 49 trio pedigrees. The distributions behind these median values are shown Supplementary Figure 11. Since we could not generate a callset with Pamir using all 147 samples at once we did not apply the Hardy-Weinberg filter. Therefore, there are no observations for Pamir in the bottom row of Supplementary Figure 10.

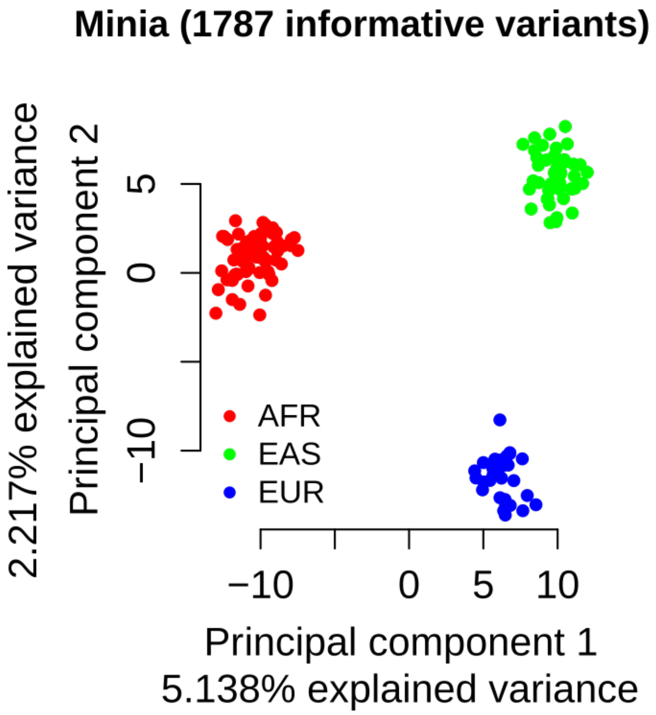


**Supplementary Figure 12.** *Principal component analysis (PCA) of non-reference sequence variants from the PDC/PKC trio pedigrees. For the PCA, the predicted genotypes of all individuals were converted into a variant/sample matrix [Niehus* et al.*, 2021] and duplicate variant as well as those with a significant Spearman correlation coefficient (P value threshold 0.05) were removed from the matrix. Finally, 1787 informative variants were used for the PCA. Each color encoded dot is an individual from one of the three continental superpopulations (AFR, African; EAS, East Asian; EUR, European).*
